# A single PA-X mutation in bovine-origin H5N1 influenza virus reduces pathogenicity in mice

**DOI:** 10.64898/2026.05.09.724031

**Authors:** Nathaniel Jackson, Mahmoud Bayoumi, Vinay Shivanna, Anna Allué-Guardia, Ashley Gay-Cobb, Ramya S. Barre, Jordi B. Torrelles, Chengjin Ye, Ahmed M. Elsayed, Luis Martinez-Sobrido

## Abstract

Dairy cows have emerged as a reservoir for human infection with highly pathogenic avian influenza (HPAI) H5N1. At the bovine–human interface, H5N1 strains may acquire adaptive mutations that influence their zoonotic potential. Sequence analysis identified a K142E substitution (bovine to human) in the PA and PA-X proteins, with the potential to affect both polymerase activity and host shutoff. Here, we used a loss-of-function approach to investigate how the bovine substitution (E142K) in PA/PA-X impacts viral replication, host shutoff activity, and pathogenicity in the human H5N1 background. Viral growth kinetics demonstrated that the virus containing the E142K substitution is attenuated, with reduced replication compared to wild-type (WT) virus. Consistently, PA-X-mediated host shutoff activity was reduced, resulting in increased induction of interferon (IFN) responses relative to WT. *In vivo*, mice infected with the E142K mutant virus survived, whereas infection with the WT virus was uniformly lethal. Despite comparable viral titers and inflammation score in mouse lungs, cytokine and chemokine profiling revealed distinct immune responses, with reduced CCL2 and increased CCL5 and IFN-γ in mice infected with the E142K mutant virus compared to mice infected with the WT virus. These findings indicate that increased virulence of the human-adapted strain is driven by a PA-X mutation that modulates inflammatory responses, producing distinct immune signatures linked to host survival or viral lethality rather than changes in polymerase activity by PA. Collectively, these results highlight PA-X as a key determinant of pathogenicity of H5N1 and a potential target for the rational design of antiviral strategies.

**IMPORTANCE:** Highly pathogenic avian influenza (HPAI) H5N1 viruses have recently expanded beyond their traditional avian hosts to infect mammals, where they are acquiring mutations associated with mammalian adaptation. These changes raise the concern that influenza H5N1 viruses could evolve the capacity for sustained human infection and human-to-human transmission and pose a pandemic threat. Therefore, it is critical to identify and functionally characterize emerging mutations that influence viral pathogenicity and host interactions. Such studies will enhance our understanding of the requirements for efficient infection and disease in mammalian hosts and inform the rational design of antiviral strategies. In this study, we present data characterizing a bovine-to-human substitution (K142E) in the viral PA-X impacting viral replication, host shutoff activity, and pathogenicity. Our results demonstrate the key role of PA-X in H5N1 viral pathogenicity and the feasibility of targeting PA-X for the rational design of antivirals to control influenza infections.

## INTRODUCTION

In March 2024, a dairy worker in Texas was confirmed to be infected with a highly pathogenic avian influenza (HPAI) H5N1 virus (1). This spillover event, involving transmission from avian to bovine hosts and subsequently to humans, identified dairy cattle as a novel intermediary reservoir and raised significant concerns given the historically high lethality of H5N1 infections in humans (2). To better understand the viral determinants required for mammalian adaptation and pathogenicity, we established reverse genetics systems for both bovine- and human-origin H5N1 strains (A/bovine/Texas/24-029328/2024 H5N1 and A/Texas/37/2024 H5N1, respectively) (3). In mice, the human strain was more pathogenic compared to the bovine strain, providing a unique opportunity to define the viral mutations that drive enhanced disease (3).

Comparative analysis between these early detected bovine and human strains in Texas revealed only nine amino acid differences in the viral PB2 (G362E, E627K, L631M), PB1 (I392V), PA (K142E, I219L, R497K), NA (S71N), and NS1 (Q40R) (3). Previous work from our laboratory demonstrated that mutations within the polymerase complex, particularly PB2 E627K and G362E, were major contributors of the increased replication and pathogenicity of the human H5N1 viral isolate (4), consistent with prior reports (5). However, another mutation of interest, PA K142E, was previously described in influenza A/Vietnam/1203/2004 H5N1 to enhance pathogenicity in mice, marked by increased viral load in organs (6). Notably, this mutation maps not only to PA but also to PA-X, a viral protein generated by ribosomal frameshifting that possesses endonuclease activity and mediates host shutoff by degrading host transcripts (7). Because PA-X was not characterized at the time of these earlier studies, the potential contribution of K142E substitution to host shutoff and pathogenesis remains unclear.

Given the dual mapping of this mutation to both PA and PA-X, we hypothesized that PA/PA-X K142E substitution contributes to viral pathogenicity not solely through effects on polymerase activity but also by modulating host shutoff. To test this hypothesis, we employed a loss-of-function strategy, investigating the effect of PA/PA-X E142K substitution (human to bovine). We first assessed polymerase activity using a minigenome (MG) assay and observed a modest, but not statistically significant, reduction associated with the mutant PA. Using reverse genetics, we then generated recombinant low pathogenic forms of influenza A/Texas/37/2024 H5N1 viruses expressing either WT (E142) or the mutant (K142) PA/PA-X and found that the mutant virus was attenuated *in vitro.* Importantly, functional assays revealed that WT virus exhibited greater host shutoff activity and more effective suppression of interferon (IFN) responses compared to the mutant virus.

*In vivo*, these differences translated into strikingly divergent outcomes: WT virus infection was uniformly lethal, whereas infection with the PA/PA-X mutant resulted in complete survival, despite comparable viral titers in the lung and nasal turbinate of infected mice. Analysis of host responses in the lungs of infected mice revealed distinct immune signatures, with WT virus infection associated with increased monocyte and macrophage-associated chemokines. In contrast, infection of mice with the mutant virus was associated with enhanced NK and T cell responses and increased IFN-γ production, suggesting that reduced PA-X-mediated host shutoff activity present in the mutant virus contributes to attenuation of the bovine-origin phenotype *in vivo*.

Finally, structural modeling provides a potential mechanistic explanation for these observations, suggesting that K142E substitution may disrupt a stabilizing interaction and alter PA-X spatial conformation and activity. Altogether, this study indicates how a single amino acid bovine-to-human substitution (K142E) in PA-X can impact viral replication, host shutoff activity and shift the balance between lethality and survival, highlighting the critical role of PA-X-mediated host shutoff in H5N1 viral pathogenesis, and the feasibility of targeting PA-X for the rational development of to control influenza infections.

## RESULTS

### PA E142K does not alter viral polymerase activity

To determine whether the PA E142K substitution affects viral polymerase activity, we utilized a minigenome (MG) assay to assess expression of a viral-like RNA in the presence of the H5N1 replication machinery (4). HEK293T cells were co-transfected with a MG plasmid encoding a dual reporter consisting of Zs-Green (ZsG) fused to Nano luciferase (Nluc), flanked by the non-coding regions (NCRs) of the viral nucleoprotein (NP) segment and driven by a human Pol I promoter (hPolI_p_) and mouse Pol I terminator (mPolI_t_) (**Fig. 1A**). In addition, plasmids encoding PB2, PB1, PA, and NP under a polymerase II cytomegalovirus (CMV) promoter were included to reconstitute the viral polymerase complex. A Cypridina luciferase (Cluc) reporter under a CAG promoter (pCAGGS plasmid) was co-transfected to normalize for transfection efficiencies (4). Polymerase activity was evaluated 30 h post-transfection by assessing both ZsG fluorescence in transfected cells and Nluc activity in cell culture supernatants (4). The following conditions were tested: (i) MG alone (MG); (ii) MG and all the polymerase components, except PA (−PA) as a negative control; (iii) PA wild-type (PA WT); and (iv) PA E142K.

**Figure 1:**
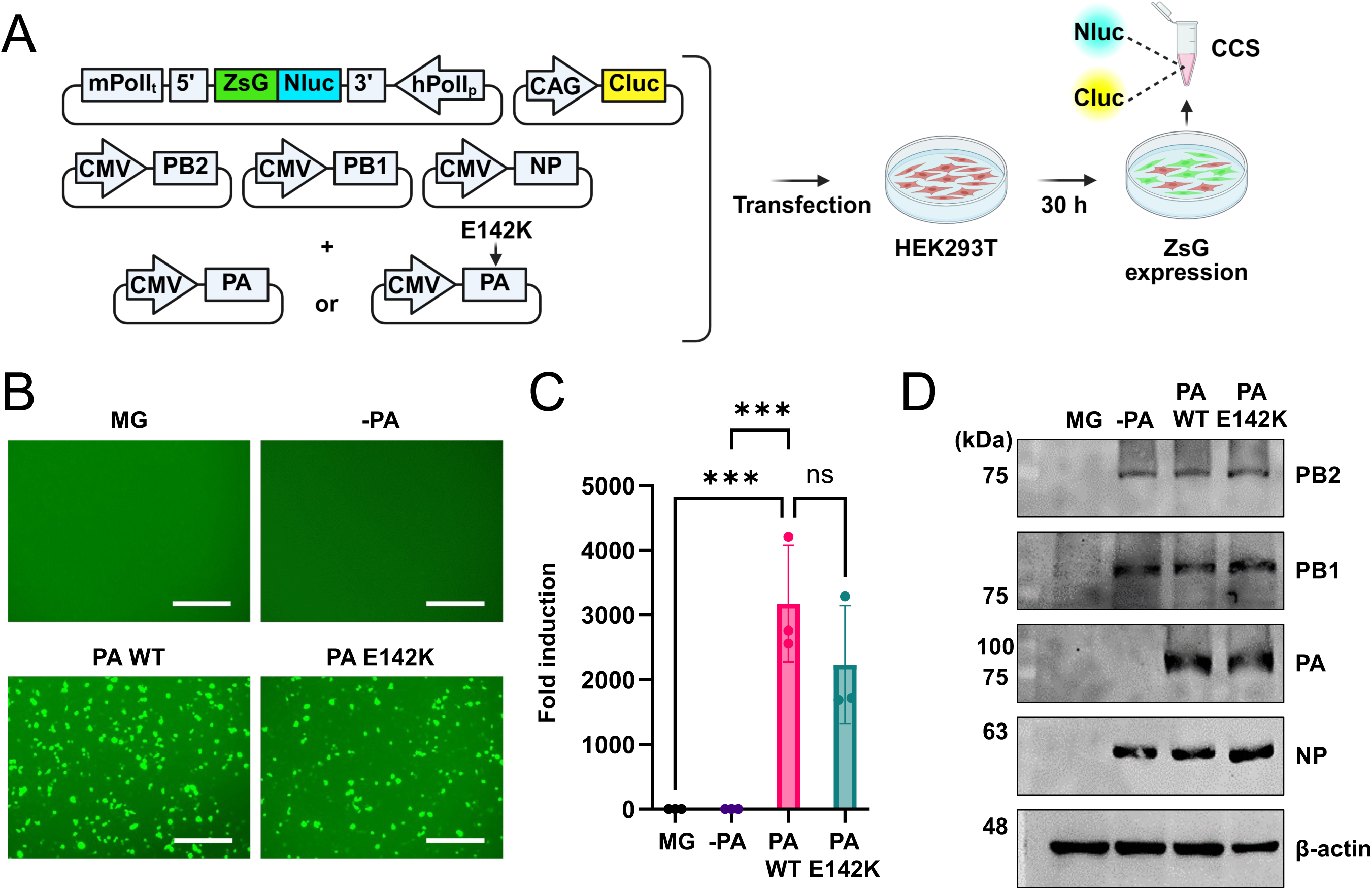
PA E142K does not alter viral polymerase activity. (**A**) Schematic representation of the MG assay. (**B-C**) HEK293T cells (6 well plate format, 1x10^6^ cells/well, triplicates) were co-transfected with plasmids encoding the viral polymerase subunits (PB2, PB1, PA) and NP under a CMV promoter, along with a MG reporter plasmid encoding ZsGreen (ZsG) fused to Nano luciferase (Nluc), flanked by the 5’ and 3’ NP non-coding regions (NCRs) under the human Pol I promoter and mouse Pol I terminator. Cypridina luciferase (Cluc), expressed from a CAG promoter, was used as a transfection control, and cells transfected in the absence of PA (–PA) were used as negative control. Wild-type (WT) PA (E142) or mutant PA (E142K) was used to assess the effect of the mutation 142 in PA on polymerase activity. At 30 h post-transfection, ZsG fluorescence was imaged by fluorescence microscopy (Scale bar = 300µm) (**B**), and Nluc and Cluc activities were measured from cell culture supernatants. Nluc activity was normalized to Cluc and expressed as fold induction relative to the –PA control (**C**). Statistical analysis was performed using an Ordinary one-way ANOVA with Dunnett’s multiple comparisons test. The data represents the mean of biological replicates (n=3), with standard deviation (SD). ns: non-significant; ***p < .001. (**D**) Cell lysates from same transfected cells were collected and analyzed by Western blot using PB2, PB1, PA, and NP antibodies to confirm expression of the viral polymerase subunits and NP, with β-actin used as a loading control.

As expected, no detectable fluorescence was observed in the MG alone and −PA transfection conditions (**Fig. 1B**), confirming minimal background activity in the absence of a functional polymerase complex (4). In contrast, cells expressing PA WT exhibited robust ZsG fluorescence expressing, indicating efficient polymerase activity. The PA E142K mutant showed a slight reduction in ZsG signal compared to WT PA. Consistent with these observations, quantification of Nluc activity (normalized to Cluc) revealed a modest decrease in fold induction for PA E142K mutant relative to PA WT; however, this difference was not statistically significant (**Fig. 1C**). Western blot analysis confirmed comparable expression levels of all the viral polymerase components across the different experimental conditions, including the PA WT and PA E142K mutant (**Fig. 1D**). Together, these results indicate that the PA E142K substitution does not significantly impair viral polymerase activity in the MG assay as compared to PA WT, despite a slight downward trend.

### PA E142K reduces viral replication

To evaluate the impact of PA E142K substitution on viral replication during authentic viral infection, we generated recombinant (r) low pathogenic avian influenza (LPAI) A/Texas/37/2024 H5N1 WT and PA E142K mutant viruses expressing monobasic HA proteins and compared their ability to replicate in different cell lines (3).

Following viral rescue, plaque assays were performed in MDCK cells to assess plaque morphology. Crystal violet staining revealed that the PA E142K mutant virus produced visibly smaller plaques compared to PA WT (**Fig. 2A**, **left**). Quantification of plaque size diameter confirmed a significant reduction in mean plaque size for the PA E142K mutant virus compared to PA WT virus (**Fig. 2A**, **right**), correlating with the reduced replication in the MG assays (**Fig. 1**). To assess replication fitness, multistep growth kinetics were performed in different mammalian cells, including MDCK (**Fig. 2B**) MDBK (**Fig. 2C**), A549 (**Fig. 2D**), and Vero-E6 (**Fig. 2E**) cells at a multiplicity of infection (MOI) of 0.001. Consistent with the plaque phenotype in MDCKs (**Fig. 2A**), the PA E142K mutant virus exhibited reduced replication compared to the virus containing PA WT across all cell types, with the most pronounced differences observed at 24 h post-infection. Notably, the magnitude of attenuation varied by cell type. The smallest reduction in viral growth was observed in MDBK (**Fig. 2C**) and Vero-E6 (**Fig. 2E**) cells, whereas the greatest reduction was observed in human A549 cells (**Fig. 2D**). Together, these findings demonstrate that PA E142K substitution reduces viral replication in mammalian cell cultures, with the strongest effect observed in human epithelial cells.

**Figure 2:**
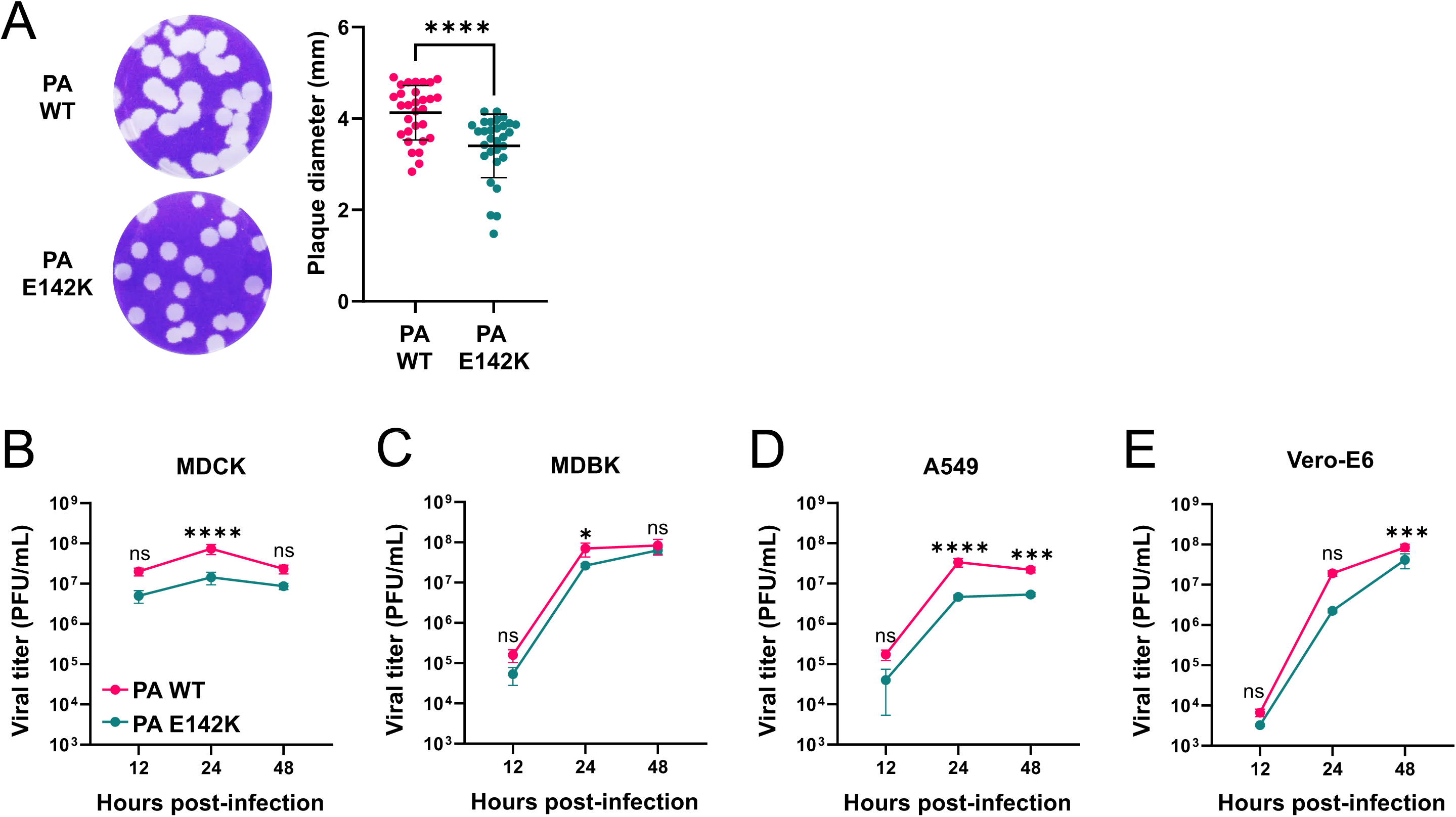
PA E142K reduces viral replication. (**A**) Plaque morphology (left) and quantification of viral plaque sizes (right) in MDCK cells infected with recombinant (r)H5N1 expressing PA wild-type (WT) (top) or PA E142K mutant (bottom). Statistical analysis was performed using a Mann–Whitney test. Data represent mean plaque diameter (n = 30) with standard deviation (SD). ****P < 0.0001. (**B-E**) Replication kinetics of rH5N1 expressing PA WT (magenta) or PA E142K substitution (teal) in MDCK (**B**), MDBK (**C**), A549 (**D**), and (**E**) Vero-E6 cells infected with MOI = 0.001. Viral titers in cell culture supernatants collected at the indicated times post-infection were determined by standard plaque assay in MDCK cells. Limit of detection (LOD) = 20 PFU. Statistical analysis was performed using a two-way ANOVA with Šídák’s multiple comparisons test. The data represents biological replicates (n=3). ns: non-significant; *p < .05, ***p < .001, ****p < .0001.

### PA E142K reduces inhibition of host gene expression in A549 cells

Because the PA segment also encodes the host shutoff protein PA-X, we next assessed whether the E142K mutation alters host shutoff activity. To evaluate temporary inhibition of global host protein synthesis, we performed a puromycin incorporation assay in IFN-deficient Vero-E6 cells, in which the two viruses showed the most similar growth kinetics, and in IFN-competent A549 cells, in which the growth difference was greatest, with PA WT virus replicating more efficiently than the PA E142K mutant (**Fig. 2**). Cells were infected with either PA WT or PA E142K mutant viruses at a MOI of 0.001 and, at 24 h post-infection, cells were incubated with puromycin (10 μg/mL) for 10 min prior to cell lysis, and protein expression evaluated by Western blot.

In Vero-E6 cells, both PA WT and E142K mutant viruses potently inhibited puromycin incorporation, as indicated by the near absence of puromycin smear compared with the strong smear observed in mock-infected cells (**Fig. 3A**, **left panel**). Notably, viral NP expression was comparable between the two viruses. In contrast, in A549 cells, mock-infected and PA E142K-infected cells showed similarly strong puromycin incorporation, whereas PA WT-infected cells showed reduced puromycin incorporation (**Fig. 3A**, **right panel**), consistent with inhibition of host protein synthesis. In A549 cells, viral NP was detected only in cells infected with the WT but not the PA E142K mutant virus, correlating with lower viral infectivity in A549 cells (**Fig. 2**). Quantification of the puromycin signal normalized to β-actin supported these observations (**Fig. 3A**, **bottom**).

**Figure 3:**
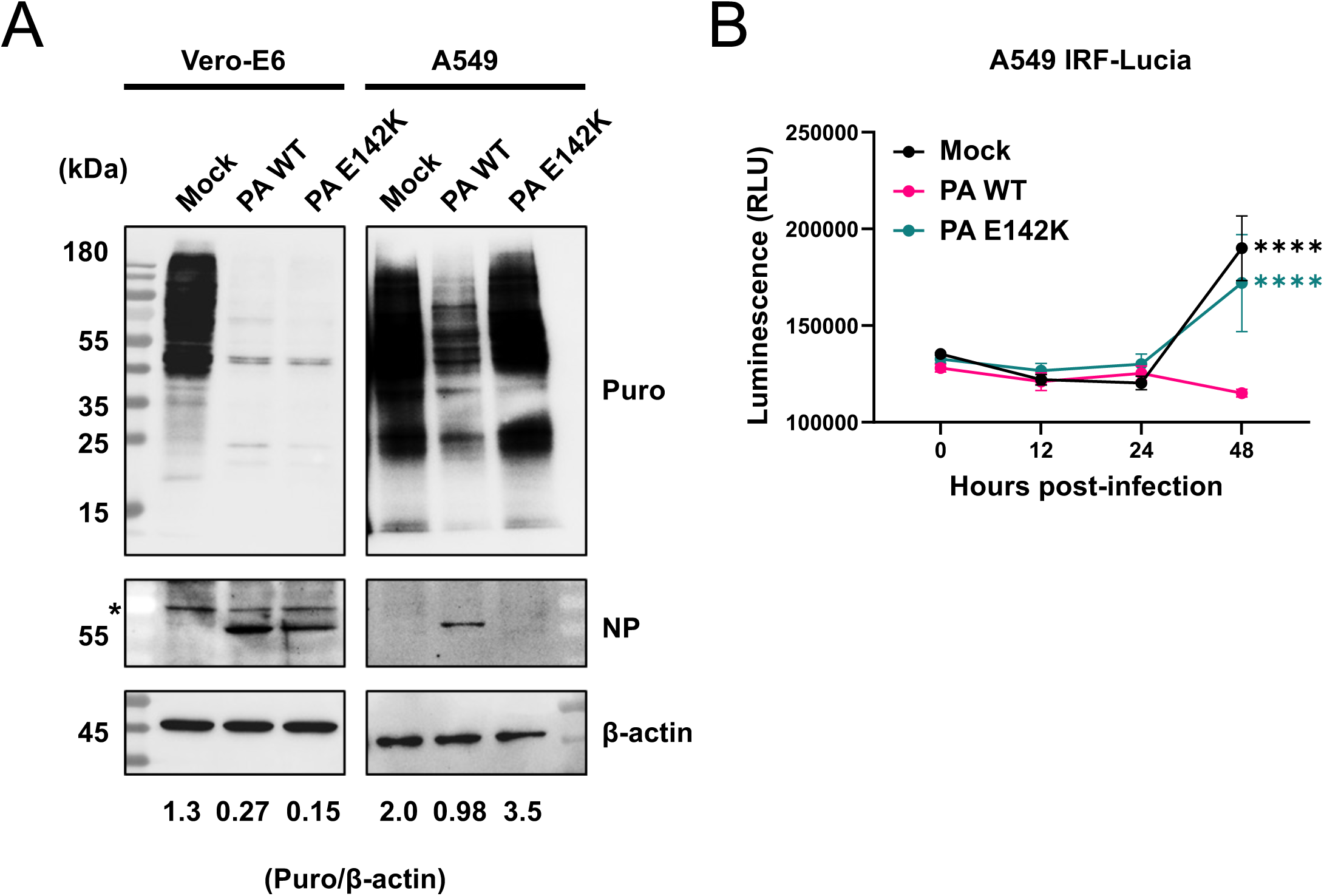
PA E142K reduces inhibition of host gene expression in A549 cells. (**A**) Vero-E6 (left) and A549 (right) cells were mock-infected or infected (MOI = 0.001) with recombinant (r)H5N1 viruses expressing PA wild-type (WT) or PA E142K. At 24 h post-infection, cell culture media were replaced with media containing puromycin (10 µg/mL) for 10 min. Cell lysates were collected and analyzed by Western blot to assess host translation via puromycin incorporation, using NP as an infection control and β-actin used as a loading control. Relative translation as determined by the ratio of puromycin to β-actin was determined and plotted for both cell lines (bottom). Non-specific bands are denoted by asterisk (*). (**C**) A549 cells expressing Lucia luciferase reporter for IFN regulatory factor (IRF) pathway induction, were mock-infected or infected (MOI = 0.001) with recombinant (r)H5N1 viruses expressing wild-type (WT) or PA E142K. Luciferase activity in cell culture supernatants collected at indicated timepoints post-infection was measured using a luciferase plate reader. Statistical analysis was performed using a two-way ANOVA with Tukey’s multiple comparisons test. The data represents the average of three biological replicates with standard deviation (SD). ****p < 0.0001.

To specifically assess IFN induction, we used A549 IFN response factor (IRF) Lucia reporter cells, which stably express a secreted Lucia luciferase reporter under the control of an ISG54 minimal promoter containing IFN-stimulated response elements (ISREs). A549 IRF-Lucia cells were infected with either PA WT or PA E142K viruses at a MOI of 0.001, and reporter activity was measured in cell culture supernatants at 12, 24, and 48 h post-infection. Mock-, PA WT-, and PA E142K-infected cells were indistinguishable through 24 h post-infection. However, by 48 h post-infection, mock-and PA E142K-infected cells showed increased reporter activity, whereas PA WT infection suppressed this response (**Fig. 3B**). Collectively, these results show that in IFN-incompetent Vero cells, host shutoff activity is comparable between PA WT and PA E142K mutant viruses. However, in IFN-competent human A549 cells, WT virus suppresses host gene expression more effectively, as demonstrated by reduced puromycin incorporation and reduced IFN reporter induction relative to PA E142K mutant-infected cells.

### PA E142K attenuates H5N1 pathogenicity *in vivo*

To assess the impact of PA E142K substitution on viral pathogenesis *in vivo*, 6-week-old female C57BL/6 mice (n=4/group) were mock-infected or infected, intranasally, with 100 PFU of PA WT or E142K mutant viruses. We have previously shown that infection with 100 PFU of PA WT is fully lethal using this mouse model of infection (3). Survival (**Fig. 4A**) and body weight (morbidity, **Fig. 4B**) were monitored daily for 15 days. All PA WT-infected mice succumbed to viral infection by 7 days post-infection, whereas all mice infected with the PA E142K mutant survived the 15-day study period. PA WT-infected mice displayed rapid weight loss beginning at 3 days post-infection, whereas PA E142K–infected mice showed a more gradual decline in body weight until day 9, followed by recovery to baseline starting at 10 days post-infection and complete recovery by day 15. To determine whether these differences in disease outcome were associated with altered viral replication, viral titers in the lungs (**Fig. 4C**) and nasal turbinate (**Fig. 4D**) in a group of mice similarly infected with PA WT and PA E142K mutant viruses (n = 4/group) were measured by plaque assay at 4 and 6 days post-infection. Lung viral titers ranged from 10⁵ to 10⁶ PFU/mL at both time points, with no significant differences observed between PA WT- and PA E142K-infected mice. Viral titers in the nasal turbinate were lower than those detected in the lungs of infected mice, reaching up to 10² PFU/mL and 10³ PFU/m at 4 and 6 days post-infection, respectively, with no significant differences between both groups. Together, these results demonstrate that PA E142K substitution attenuates the pathogenicity of H5N1 in mice despite comparable levels of viral replication in the upper and lower respiratory tract.

**Figure 4:**
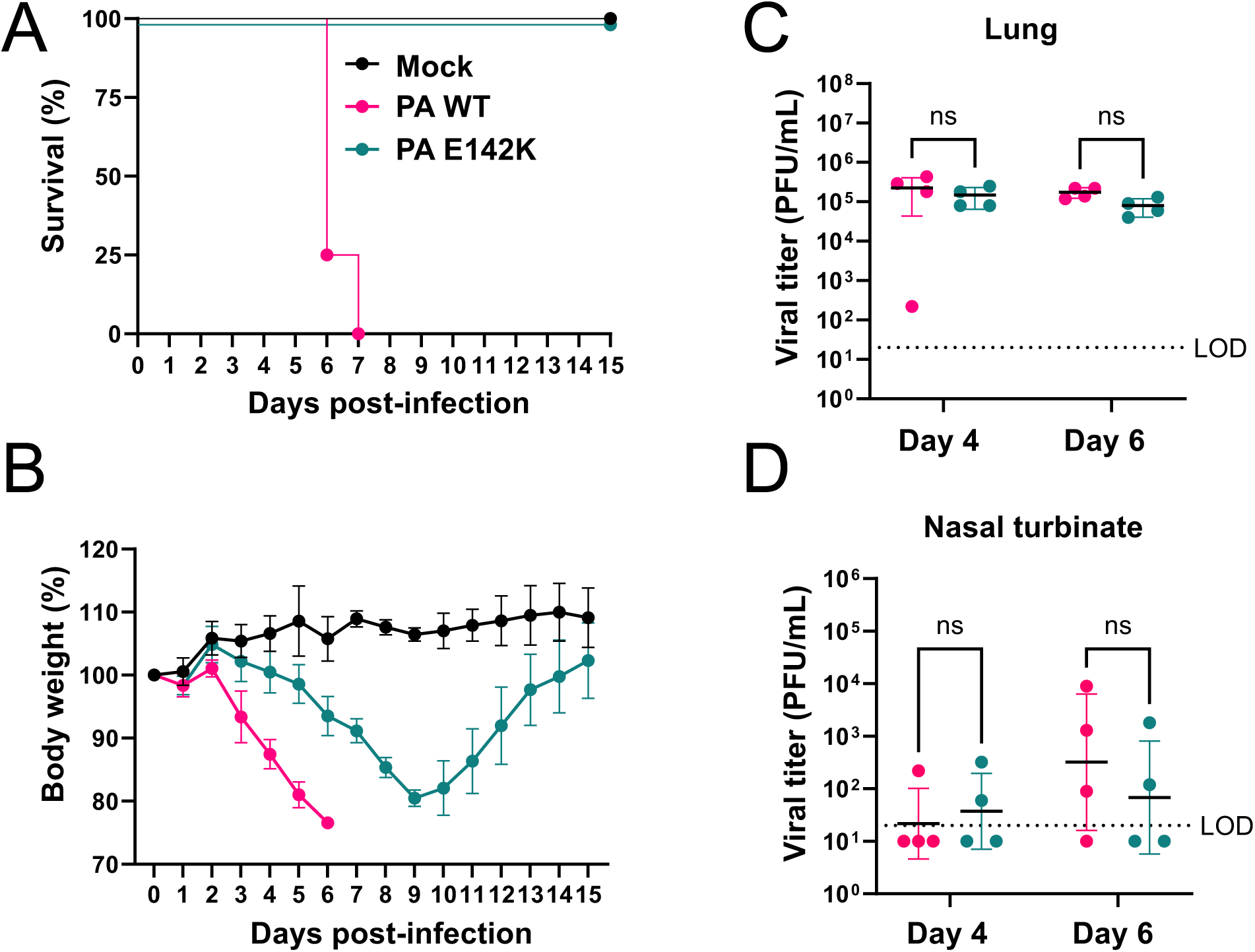
PA E142K attenuates H5N1 *in vivo*. **(A-B)** Six-week-old female C57BL/6 mice (n=4) were mock-infected or infected intranasally with 10² PFU of recombinant (r)H5N1 expressing PA WT or PA E142K. Mice were monitored daily for survival (**A**) and body weight (**B**). (**C & D**) Lungs and nasal turbinate from mice infected intranasally with 10² PFU of recombinant (r)H5N1 expressing WT or PA E142K (n = 4 per group) were collected at 4- and 6-days post-infection for viral load determination by standard plaque assay in MDCK cells. Limit of detection (LOD) = 20 PFU. Values below LOD are plotted as 10 PFU. Statistical analysis was performed using a two-way ANOVA with Šídák’s multiple comparisons test. ns: non-significant.

### PA E142K alters lung cytokine and chemokine responses

To evaluate differences in host immune responses *in vivo*, cytokine and chemokine expression in lung tissues collected at 4 and 6 days post-infection was assessed using a Luminex-based multiplex assay. Log_2_ fold-change (FC) analysis of PA E142K-infected mice relative to PA WT-infected mice revealed differential modulation of several immune markers (**Fig. 5A**). At day 4 post-infection, lungs from mice infected with the PA E142K mutant exhibited downregulation of pro-inflammatory cytokines, including IL-6, TNF, and IFN-β, as well as chemokines such as CCL2 and IP-10, which are broadly associated with recruitment of monocytes and activated immune cells (8). In contrast, CCL5, a chemokine involved in recruitment of lymphocytes and other immune cell populations (9), was upregulated. At day 6 post-infection, IL-6 remained downregulated, while IFN responses were altered, with increased expression of IFN-α and IFN-γ in PA E142K-infected mice relative to PA WT-infected mice. Statistical analysis identified CCL2 and CCL5 at 4 days post-infection and IFN-γ at 6 days post-infection, as the most significantly modulated factors (**Fig. 5B**). Individual expression profiles for these cytokines and chemokines are shown in **Figs. 5C–E**. To further characterize immune responses in the lungs of mock- and virus-infected animals, we performed histopathological analysis. Although mice infected with the PA WT virus showed a trend toward increased percent pathology at day 6 post-infection, this did not reach statistical significance (**Supplemental Fig. 1A–B**). Immunohistochemical analysis of lung sections for monocytes (CD163^+^) and T cell subsets (CD4^+^ and CD8^+^) revealed no significant differences in monocyte abundance and only modest changes in T cell populations.

**Figure 5:**
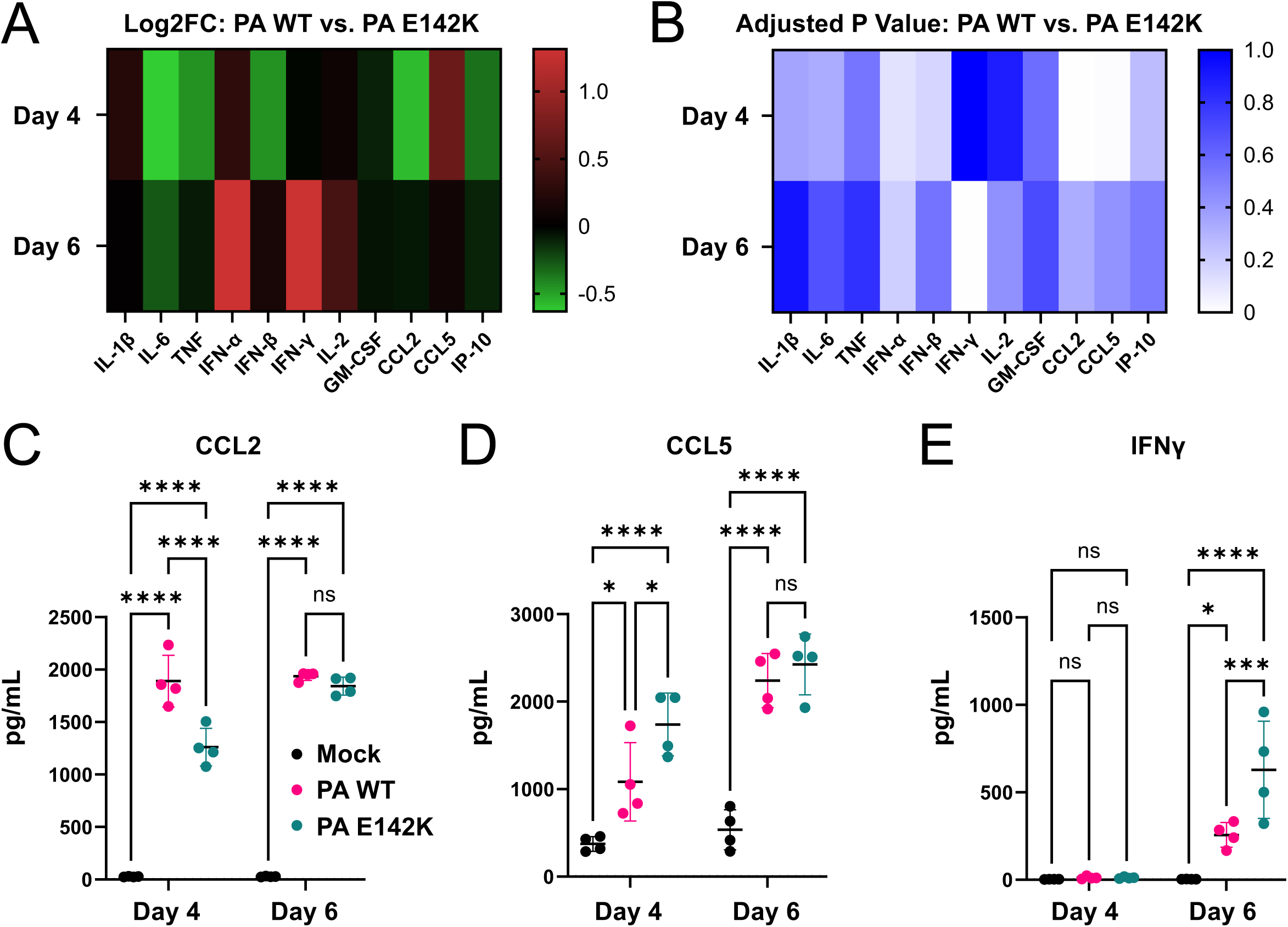
PA E142K alters lung cytokine and chemokine responses. (**A-B)** Cytokine and chemokine expression in lung homogenates from mock-infected or virus-infected mice collected at 4- and 6-days post-infection above were measured using a Luminex-based multiplex assay. Expression was analyzed as log_2_ fold change (**A**) and adjusted P value comparing recombinant (r)H5N1 expressing WT to PA E142K (**B**). (**C-E**) Proteins significantly modulated based on adjusted P value are shown for CCL2 (**C**), CCL5(**D**), and IFN-γ (**E**). Statistical analysis was performed using a two-way ANOVA with Tukey’s multiple comparisons test. Data represent mean values from mock-infected or infected animals (n=4) with standard deviation (SD). ns: not significant; *p < 0.05; ***p < 0.001; ****p < 0.0001.

CD4^+^ T cells showed a non-significant increase, whereas CD8^+^ T cells were significantly elevated in PA WT-infected mice compared to those infected with the PA E142K mutant virus at day 6 post-infection (**Supplemental Fig. 1C**). Collectively, these data indicate that infection with the PA E142K mutant virus elicits a distinct pulmonary immune profile relative to PA WT virus infection that is associated with improved survival, despite minimal differences in lung pathology and immune cell infiltration.

### Predicted structural impact of PA-X E142K

To evaluate the potential structural impact of E142K mutation on PA-X, predicted protein structures were generated using AlphaFold3 (**Fig. 6**) (10). Models corresponding to the human PA-X (E142; magenta) and the bovine (K142; teal) were aligned to assess local structural differences. Structural comparison revealed that the K142 residue is positioned to form a potential electrostatic interaction with E101 residue, with an inter-residue distance of approximately 2.720 Å, consistent with a stabilizing interaction. In contrast, the E142 substitution introduces a negatively charged side chain in proximity to E101, which may result in electrostatic repulsion and disruption of this interaction.

**Figure 6:**
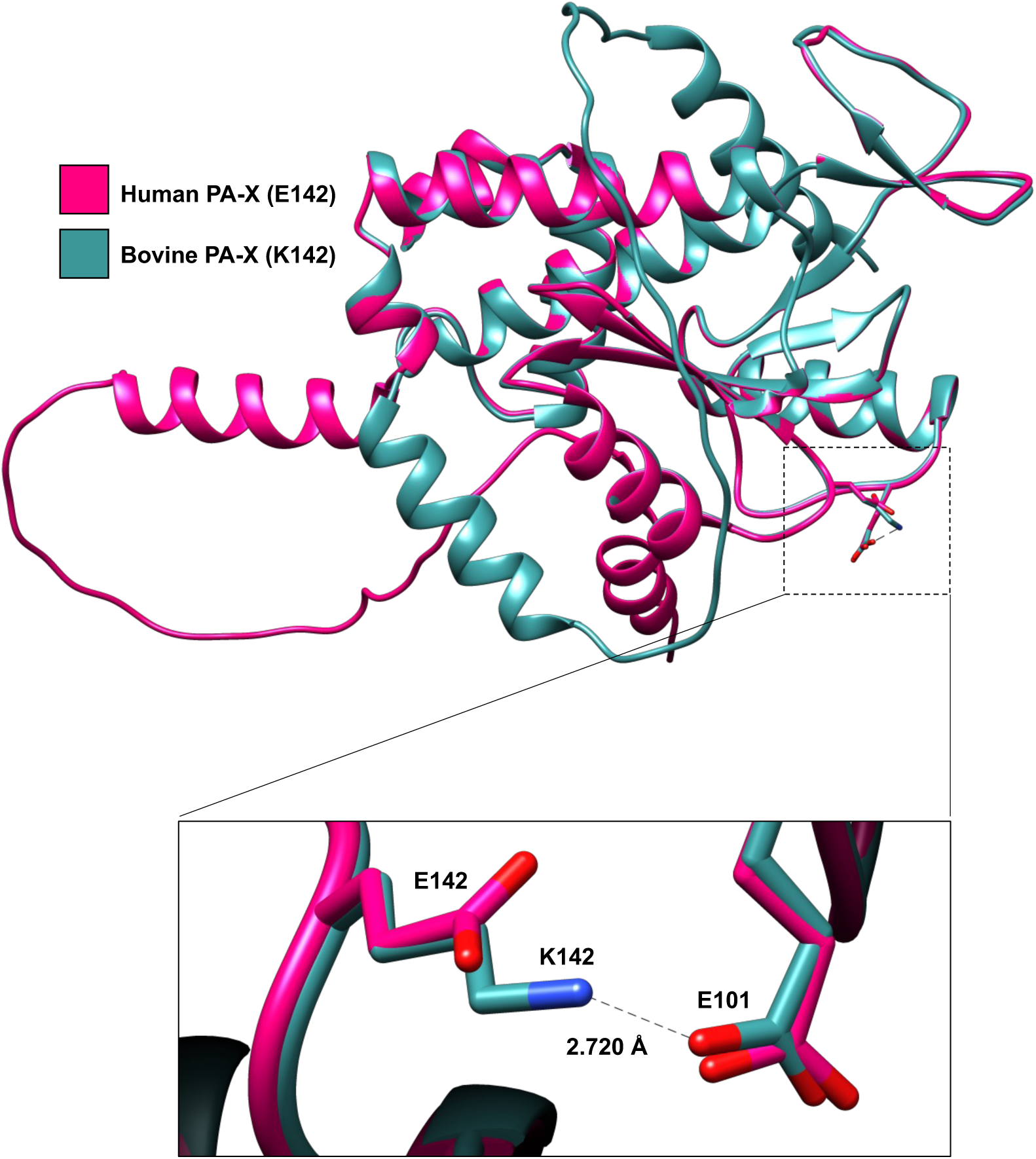
Structural basis of PA-X E142K. Predicted 3D structures of human (magenta) and bovine (teal) PA-X were generated using AlphaFold3 and visualized in Chimera. The distance between the side chains of K142 and E101 (bovine) was measured at 2.72 Å.

These observations suggest that K142 may locally stabilize the PA-X structure relative to E142, providing a potential structural basis for differences in PA-X function.

## DISCUSSION

Previous work examining an avian to human K142E substitution in influenza A/Vietnam/1203/2004 H5N1 PA segment demonstrated that E142 enhances viral replication in mouse organs compared to K142 (6). In contrast, in this study using bovine-origin human and bovine H5N1 strains, we did not observe enhanced polymerase activity of E142 (human) over K142 (bovine) using a MG reporter assay or *in vivo*, using recombinant viruses, although we did observe enhanced viral replication in cultured cells in all tested cell lines. Notably, residue 142 maps not only in PA but also in PA-X, a viral endonuclease with host shutoff activity (7) that was not characterized in the original study with influenza A/Vietnam/1203/2004 H5N1 (6).

Because polymerase activity was not significantly altered by the E142K mutation, yet differences in viral replication were observed, we hypothesized that these phenotypes may instead be driven by differences in host shutoff activity by PA-X.

In IFN-competent A549 cells (11), PA WT virus replicated more efficiently than the PA E142K mutant, whereas in IFN-incompetent Vero-E6 cells (12), replication of both viruses was more comparable, with only a modest advantage for WT over the E142K mutant virus. Consistent with this, host shutoff activity was similar between PA WT and PA E142K in Vero-E6 cells but was more pronounced for WT virus in A549 cells. These findings suggest that in A549 cells, IFN responses restrict replication of the PA E142K mutant virus, whereas PA WT virus can better evade this restriction through more effective host shutoff activity. In contrast, in IFN-deficient Vero-E6 cells, both viruses replicate similarly and differences in host shutoff activity are less apparent. This interpretation is further supported by experiments using A549 IFN response factor (IRF) Lucia reporter cells, which demonstrated increased IRF activation in PA E142K-infected cells compared to PA WT. Together, these data indicate that the E142 residue enhances PA-X-mediated suppression of host activity, including IFN responses, thereby promoting viral replication in IFN-competent environments, whereas the K142 mutation is less effective at suppressing host responses, including IFN antiviral activities, resulting in reduced replication under these conditions.

Although PA WT and E142K viruses reached comparable titers in the nasal turbinate and lungs of infected mice, their divergent clinical outcomes indicate that pathogenesis was governed primarily by host responses rather than replication levels. PA WT infection was associated with an inflammatory profile characterized by IFN-β (13–15), TNF (13, 16), IL-6 (13, 17), CCL2 (13, 14, 16), and IP-10 (14) together with increased CD8+ T-cell accumulation (16, 18), consistent with dysregulated immune responses. In contrast, mice infected with the PA E142K mutant virus exhibited reduced CCL2 and increased CCL5 (19–21) at day 4 post-infection, followed by elevated IFN-γ (20, 22, 23) at day 6 post-infection, together with increased IL-2 (20, 22) responses, consistent with a more effective antiviral program, with the caveat of elevated IFN-α, which can mediate disease severity (15). CCL2 is a key chemoattractant for monocytes and macrophages (8), whereas CCL5 recruits NK and T cells (9), which are major producers of IFN-γ and contribute to viral clearance (24, 25). Collectively, these findings suggest that the PA E142K mutation attenuates virulence by altering immune cell recruitment and modulating host inflammatory responses, favoring NK- and T-cell–associated antiviral immunity while limiting immunopathology associated with monocyte and macrophage infiltration. We therefore propose that the PA E142K mutant, which exhibits reduced host shutoff activity, causes a rebalanced immune signature with preserved early IL-1β and GM-CSF induction, but attenuated pro-inflammatory and type I IFN-associated myeloid chemokines, with increased IFN- and lymphocyte-associated mediators. In contrast, PA WT virus, with more potent host shutoff activity, likely drives a dysregulated immune response characterized by excessive monocyte/macrophage recruitment and impaired antiviral effector responses. This correlates with previous studies describing that enhanced shutoff activity of PA-X correlates with enhance morbidity and mortality (26–28).To further evaluate this hypothesis, we extended our analysis of formalin-fixed lung tissues collected at days 4 and 6 post-infection. Histopathological assessment revealed no significant differences in lung pathology between PA WT- and mutant-infected mice at either time point. One possible explanation is that pathological changes may occur earlier than our earliest timepoint of 4 days post-infection as prior studies of PA-X-deficient viruses with abrogated host shutoff activity have reported differences in pathology only 1 day post-infection which is resolved by day 2 post-infection (29). We hypothesize that PA WT-infected animals may have an earlier onset of pathology and subsequently increase the duration of acute long injury leading to uniform lethality whereas in PA mutant E142K-infected mice, pathology onset may be delayed, leading to eventual recovery due to less total time with acute lung injury.

We next quantified immune cell populations by immunohistochemistry. M2-like macrophage (CD163^+^) abundance did not differ between infected and mock-treated animals, suggesting that 4 days post-infection is likely out of range of M2 cell infiltration (30). In contrast, T cell analysis revealed increased CD4^+^ and significantly elevated CD8^+^ T cells in WT-infected mice at 6 days post-infection. This finding was unexpected given the higher levels of IFN-γ observed in PA mutant E142K-infected mice at this time point, which are typically associated with protective immune responses (22, 25). One possible explanation is that T cell functionality, rather than abundance, underlies these differences (22, 31). Consistent with this, previous studies have shown that H5N1 infection can induce functionally impaired CD8^+^ T cells characterized by increased expression of inhibitory markers such as PD-1 (18). Thus, infection with PA WT virus may drive greater T cell accumulation with reduced functional capacity compared to infection with the PA E142K mutant virus. While our data support a role for differential immune responses in the observed survival phenotype, this study is limited by the absence of early time point analysis (e.g., 1-day post-infection), which may have missed key innate immune events (32). Additionally, NK cells were not evaluated and could represent a significant source of IFN-γ (25). Future work incorporating earlier time points and comprehensive immune profiling will be necessary to fully resolve the temporal and cellular dynamics underlying these responses.

Finally, structural modeling provides a potential mechanistic basis for the observed attenuation of the PA-X E142K mutant. Specifically, K142 is predicted to interact with residue E101, potentially stabilizing a more constrained conformation of PA-X that may limit endonuclease activity. In contrast, E142 may introduce electrostatic repulsion with E101, promoting a more flexible conformation that enhances endonuclease activity and, consequently, host shutoff. However, this remains speculative and will need to be addressed experimentally. Future biochemical analyses using recombinant PA-X WT and PA-X E142K mutant proteins directly assess how the E142K substitution impacts endonuclease activity and host shutoff function.

In summary, this study demonstrates that a single mutation in H5N1 PA segment can determine survival versus lethality *in vivo*. Although PA is a critical component of the viral polymerase complex and would be expected to primarily influence viral replication, our findings indicate that this mutation, also mapping to the PA-X host shutoff protein, plays a more prominent role in modulating host shutoff activity than polymerase function. Importantly, despite minimal differences in viral replication *in vivo*, the mutation profoundly altered disease outcome, underscoring the central role of host response modulation in H5N1 pathogenesis. These results highlight PA-X-mediated host shutoff as a key determinant of virulence and suggest that targeting this function may represent a promising strategy for the rational development of antivirals. More broadly, this work provides insight into mechanisms driving the pathogenicity of the recently emerged bovine-origin H5N1 influenza A virus and informs efforts to mitigate its ongoing public health threat.

## MATERIALS AND METHODS

### Biosafety

All *in vitro* and *in vivo* experiments with low pathogenic avian influenza (LPAI) H5N1 viruses were performed in biosafety level 3 (BSL-3) and animal BSL-3 (ABSL-3) facilities, respectively, at Texas Biomedical Research Institute (Texas Biomed). All procedures were approved by the Institutional Biosafety Committee (IBC) and Institutional Animal Care and Use Committee (IACUC) at Texas Biomed (21-017 and 1785MU, respectively).

### Antibodies

Monoclonal antibodies against H5N1 polymerase proteins were obtained from BEI Resources, including 170-3C12 (PB2), F5-46 (PB1), and 1F6 (PA). Viral NP was detected using a mouse monoclonal antibody, HT103 (Kerafast), as previously described (4). β-actin was detected using the mouse monoclonal antibody AC-15 (Sigma). For the puromycin incorporation assay, NP and β-actin were detected using the same HT103 and AC-15 mouse monoclonal antibodies, and puromycin-labeled proteins were detected using the mouse anti-puromycin antibody 12D10 (Sigma).

### Cell lines

Human embryonic kidney (HEK293T, ATCC CRL-3216), Madin-Darby kidney (MDCK, ATCC CCL-34), human lung adenocarcinoma epithelial (A549, ATCC CCL-185), African green monkey kidney epithelial cells (Vero-E6; ATCC CRL-1586), MDBK cells (a gift from Dr. Daniel Perez), and A549 cells expressing Lucia gene under the control of IFN-stimulated response elements (A549-IRF-Lucia, InvivoGen a549d-nfis) were maintained in cell culture medium constituted of Dulbecco’s modified eagle medium (DMEM) (Corning) supplemented with 5-10% fetal bovine serum (FBS) (VWR) and 100 units/mL penicillin streptomycin L-glutamine (Corning), at 37°C in a 5% CO_2_ incubator.

### Viruses and plasmids

Recombinant LPAI A/Texas/37/2024 H5N1 (rLPhTX) WT virus containing a monobasic cleavage site in the viral HA was previously generated and described (3). To generate the rLPhTX expressing PA E142K substitution, the pHW2000 plasmid encoding PA was mutated from E to K by site-directed mutagenesis. Rescue of rLPhTX PA E142K was performed as previously described for rLPhTX WT (3). Briefly, the eight pHW2000 plasmids required for viral rescue were co-transfected into a co-culture of HEK293T and MDCK cells using Lipofectamine 3000 (Thermo Fisher Scientific) according to the manufacturer’s instructions. The transfection mixture was replaced with cell culture medium containing 0.2% bovine serum albumin (BSA) (Sigma) and 2 μg/mL TPCK-treated trypsin (Sigma). At 72 h post-transfection, cell culture supernatants were collected, clarified by centrifugation, and used to infect fresh MDCK cells in T75 flasks to generate viral stocks in medium containing 0.2% BSA and 1 μg/mL TPCK-treated trypsin. At 72 h post-infection, when 100% cytopathic effect was observed, cell culture supernatants were clarified and aliquoted.

### Minigenome (MG) assays

The MG plasmid encoding Zs-Green (ZsG) fused to Nano luciferase (Nluc), flanked by the 5′ and 3′ non-coding regions (NCRs) of the H5N1 NP segment and driven by a human Pol I promoter and mouse Pol I terminator, as well as pHW2000 plasmids encoding the viral polymerase subunits (PB2, PB1, PA) and NP (expressed under a CMV promoter), were previously generated (4). Cypridina luciferase (Cluc) expressed from a pCAGGS vector (CAG promoter) was used as a transfection control. The MG assay was performed as previously described (4). Briefly, HEK293T cells (12-well format, 5x10⁵ cells/well, triplicates) were co-transfected with a MG plasmid encoding ZsG-Nluc along with plasmids encoding the viral polymerase proteins (PB2, PB1, PA) and NP under a CMV promoter, and the pCAGGS Cluc. Plasmids were co-transfected using Lipofectamine 3000 (Thermo Fisher Scientific) according to the manufacturer’s instructions. Co-transfections in the absence of the plasmid encoding PA (-PA) were included as negative controls. At 6 h post-transfection, the transfection mixture was replaced with fresh cell culture medium. At 30 h post-transfection, Nluc and Cluc activities were measured using luciferase assay kits (Promega and Thermo Fisher Scientific, respectively) according to the manufacturers’ instructions and quantified using a luciferase plate reader (Promega). Nluc fold induction was calculated by dividing the Nluc activity by the Cluc activity for each sample and expressed as fold change relative to the negative control group lacking PA. ZsG fluorescence was imaged using a fluorescence microscope (Thermo Fisher Scientific). After imaging, cell monolayers were washed in PBS and lysed in NP-40 lysis buffer (Thermo Fisher Scientific) for Western blot analysis.

### Western blot

Western blotting was performed as previously described (4). Briefly, cell lysates prepared in NP-40 lysis buffer were mixed with SDS-PAGE loading buffer, boiled, resolved by SDS-PAGE (12%), and transferred to nitrocellulose membranes (Bio-Rad). Membranes were blocked in 5% nonfat dry milk in PBS containing Tween-20 (PBST) for 1 h, followed by overnight incubation with primary antibodies. Membranes were washed and incubated with horseradish peroxidase (HRP)-conjugated secondary antibodies (Sigma) for 1 h, washed, and developed using a chemiluminescence substrate kit (Thermo Fisher Scientific) according to the manufacturer’s instructions.

Chemiluminescence was imaged using a ChemiDoc imaging system (Bio-Rad).

### Virus titration

MDCK cells (6-well plate format, 1x10^6^ cells/well) were infected with 10-fold serial dilutions of virus in cell culture media containing 0.2% BSA. After 1 h, the viral inoculum was removed and replaced with 4 mL of semi-solid overlay containing DMEM/F12 (Gibco), 100 units/mL penicillin streptomycin L-glutamine (Corning), 0.2% BSA (Sigma), 10mM HEPES (Gibco), 0.2% NaHCO_3_ (Sigma), 0.01% DEAE-dextran (MP Biomedicals), 1.25% Avicel (IMCD), and 1 μg/mL TPCK-treated trypsin (Sigma). At 48-72 h post-infection, semi-solid overlay was removed and replaced with 10% neutral buffered formalin (VWR) for overnight inactivation. Formalin was removed and cell monolayers were stained with 0.2% crystal violet (Fisher Scientific) for 5 min. Plates containing crystal violet plaques were imaged with a scanner (Epson) and plaque diameter was measured using ImageJ for 30 individual plaques per virus tested.

### Viral growth kinetics

MDCK, MDBK, A549, and Vero-E6 cells (6-well plate format, 1x10^6^ cells/well, triplicates) were infected with rLPhTX viruses (WT or PA E142K) using a MOI of 0.001. After 1 h, the viral inoculum was removed and replaced with cell culture media containing 0.2% BSA and TPCK-treated trypsin (Sigma) at 1 μg/mL for MDCK, MDBK, and Vero-E6 cells and 0.25 μg/mL for A549 cells. At 12, 24, and 48 h post-infection, cell culture supernatants were collected, and viral titers were determined by standard plaque assay in MDCK cells (12-well plate format, 5x10⁵ cells/well, duplicates).

### Puromycin incorporation assay

A549 cells (6-well plate format, 1x10⁶ cells/well, triplicates) were mock-infected or infected with rLPhTX (WT or PA E142K) using a MOI of 0.001. After 1 h, the viral inoculum was removed and replaced with cell culture medium containing 0.2% BSA and TPCK-treated trypsin (Sigma) at 0.25 μg/mL. At 24 h, cell culture media was removed and replaced with media containing 10 μg/mL of puromycin (InvivoGen) for 10 min (33). Cells were washed with PBS and lysed in NP-40 lysis buffer (Thermo Fisher Scientific) for Western blot analysis.

### A549 IRF Lucia reporter assay

A549 IRF Lucia cells (6-well plate format, 1x10⁶ cells/well, triplicates) were mock-infected or infected with rLPhTX (WT or PA E142K) using a MOI of 0.001. After 1 h, the viral inoculum was removed and replaced with cell culture media containing 0.2% BSA and TPCK-treated trypsin (Sigma) at 0.25 μg/mL. At 12, 24, and 48 h post-infection, cell culture supernatants were collected and assayed for Lucia luciferase activity using QUANTI-Luc 4 reagent (InvivoGen). Lucia activity was measured by luminescence using a GloMax plate reader (Promega).

### Mouse experiments

6-week-old female C57BL/6 mice (The Jackson Laboratory) were intranasally mock-infected or infected with 100 PFU per animal of rLPhTX (WT or PA E142K). Mice were monitored daily for body weight and survival (n=4). Mice exceeding 25% body weight loss were humanely euthanized. At 2- and 4-days post-infection, mice (n=4) were humanely euthanized and lungs and nasal turbinate were collected. The left lobe of the lung was fixed in 10% neutral buffered formalin (VWR) for histopathology analysis and the remaining right lung lobes and nasal turbinate were homogenized and clarified by centrifugation for viral load determination by standard plaque assay in MDCK cells (12-well plate format, 5x10⁵ cells/well, duplicates).

### Lung cytokine and chemokine profiling

Cytokine and chemokine levels (IL-1β, IL-6, TNF, IFN-α, IFN-β, IFN-γ, IL-2, GM-CSF, CCL2, CCL5, and IP-10) were measured in lung homogenates using a custom multiplex ProcartaPlex assay (Thermo Fisher Scientific) according to the manufacturer’s instructions and as previously described (3). Data were analyzed using xPONENT software.

### Lung histopathology and immunohistochemistry

The left lung lobe was fixed in 10% neutral buffered formalin, transferred to 75% ethanol, paraffin-embedded, sectioned (4 µm), and stained with H&E as previously described (3, 4). Stained sections were evaluated by a board-certified veterinary pathologist in a blinded manner. Whole-slide images were acquired at 20× magnification (Axio Scan Z1, Zeiss) and analyzed using HALO software (Indica Labs) to quantify pathology. Immunohistochemistry was performed on formalin-fixed lung sections to assess macrophages and T cell populations (CD163, CD4, and CD8). Total cell counts positive for antibody staining were normalized to tissue area and analyzed using HALO software (Indica Labs).

### Structural modeling

The PA-X coding sequences (CDS) from A/Texas/37/2024 H5N1 (GenBank: PP577942.1) and A/bovine/Texas/24-029328-02/2024 (GenBank: PP599472.1) isolates were used for structure prediction using AlphaFold3. Predicted structures were visualized and analyzed in UCSF Chimera.

### Statistical analysis

All statistical analyses were performed using GraphPad Prism. Data are presented as mean ± standard deviation (SD). For MG assays (n = 3), statistical significance was determined using one-way ANOVA with Dunnett’s multiple comparisons test. Plaque diameter analysis (n = 30 plaques per group) was performed using a Mann–Whitney test. Growth kinetics experiments (n = 3) were analyzed using two-way ANOVA with Sidak’s multiple comparisons test. Lucia luciferase assays (n = 3) were analyzed using two-way ANOVA with Tukey’s multiple comparisons test. For in vivo studies (n = 4 mice per group), viral titers were analyzed using two-way ANOVA with Sidak’s multiple comparisons test. Pathology scores (n = 4 mice per group) were also analyzed using two-way ANOVA with Sidak’s multiple comparisons test. Cytokine and chemokine and lung immunohistochemistry analyses (n = 4 mice per group) were analyzed using two-way ANOVA with Tukey’s multiple comparisons test. Statistical significance was defined as p < 0.05. Significance levels are indicated as follows: p < 0.05 (*), p < 0.01 (**), p < 0.001 (***), and p < 0.0001 (****).

## ACKNOWLEDGMENTS

This work was supported by a grant from the American Lung Association (ALA) to L.M-S, a Texas Biomed Forum Award to A.M., and a Douglass Award to R.S.B. We thank BEI Resources for providing the PB2, PB1, and PA antibodies used in this study. We also thank Dr. Daniel Perez for providing the MDBK cells.

## SUPPLEMENTAL MATERIAL

**Supplemental Figure 1: Lung histopathology and immune cell infiltration.** (**A**) Lung sections (n=4) of mock-infected or virus-infected mice collected at 4- and 6-days post-infection, were stained by hematoxylin and eosin (H&E) and scanned. Sections were analyzed using HALO software to calculate percentage pathology in a blinded manner. Statistical analysis was performed using a two-way ANOVA with Šídák’s multiple comparisons test. The data represents the mean of lung pathology from mock-infected or virus-infected animals (n=4). ns: non-significant. (**B**) Representative day 4 and 6 lung images of mock-infected and recombinant (r)H5N1 expressing WT to PA E142K infected mice (Scale Bar = 100µm). (**C**) Cells positive for CD163, CD4, and CD8 as determined by immunohistochemistry analysis of lung sections on day 4 and 6. Sections were analyzed using HALO software to determine positive cells normalized to tissue area. Statistical analysis was performed using a two-way ANOVA with Tukey’s multiple comparisons test. Data represent mean values from mock-infected or infected animals (n=4 per group) with standard deviation (SD). ns: not significant; *p < 0.05; **p < 0.01; ***p < 0.001.

**Supplemental Figure 2: Graphical abstract.** Human A/Texas/37/2024 and A/bovine/Texas/24-029328-02/2024 H5N1 influenza strains differ by 10 amino acids across multiple viral proteins. A single mutation in the PA-X host shutoff protein (E142K) uniquely alters cytokine and chemokine responses and, subsequently, immune cell programming through differential inhibition of host gene expression, with the human PA-X (E142) more efficiently inhibiting host gene expression than the bovine PA-X (K142).

In mice, this results in viral attenuation of the bovine strain compared to the human strain.

