## Supplementary figures and images for "A single PA-X mutation in bovine-origin H5N1 influenza virus reduces pathogenicity in mice"

### Supplemental Figures

Supplemental Fig. 1

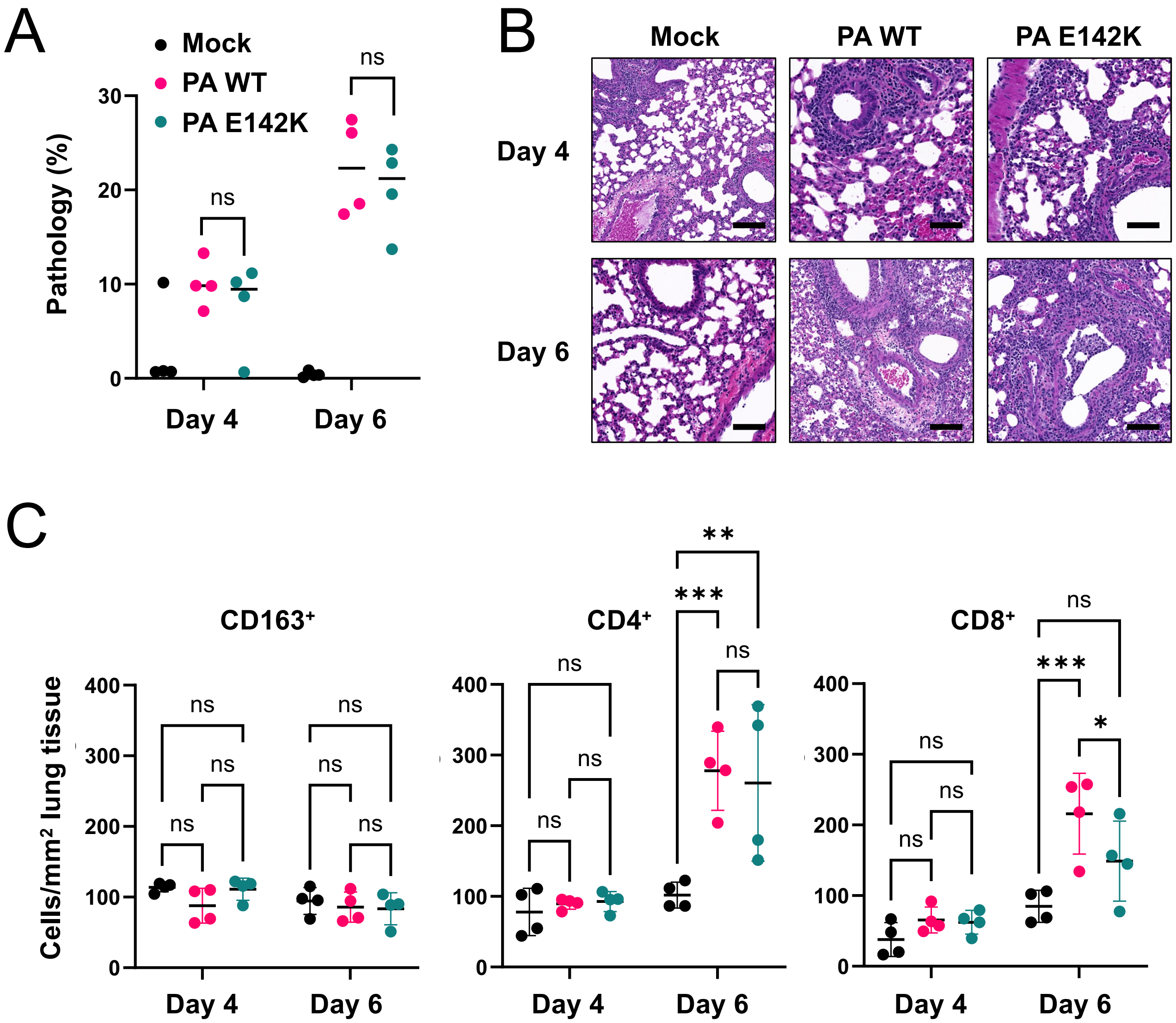

Supplemental Fig. 2

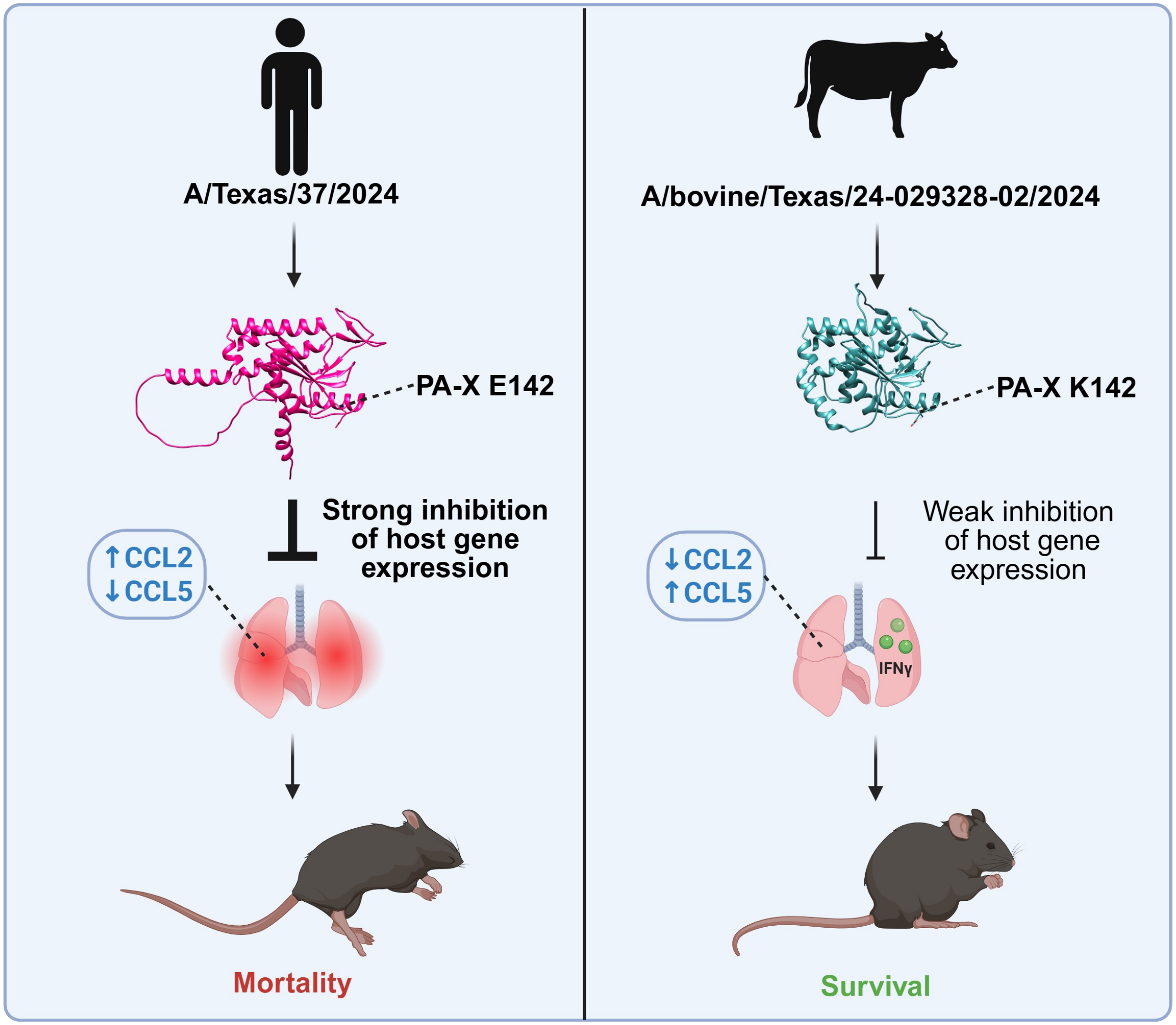
